# Alginate Nanohydroxyapatite Composite for Dental Tissue Engineering Application

**DOI:** 10.64898/2025.12.11.693830

**Authors:** Ananya N. Nayak, Roshni Ramachandran, Hansel P. Shah, Ishani Chowdhary, Punya Sharma

## Abstract

Traumatic dental tissue injury is a common health concern globally, with more than a billion people affected worldwide. In this study, we have synthesized green route nanohydroxyapatite utilizing green tea extract, which further enhances the osteogenesis, osteointegration, and biocompatibility of the synthesized hydroxyapatite. This Nanohydroxyapatite was capped with an antibiotic, integrated into an alginate scaffold, and coated with a painkiller to create a dual drug release system, providing rapid release of painkiller and sustained release of antibacterial drug. Biocompatibility and osteoblast proliferation were confirmed through MTT assays, and osteogenesis by ALP activity, validating the suitability of the scaffold for dental bone tissue engineering applications. The developed nanocomposite scaffold shows significant promise in enhancing dental bone repair by simultaneously addressing the inflammation, infection, and regeneration challenges.

**Graphical Abstract:** 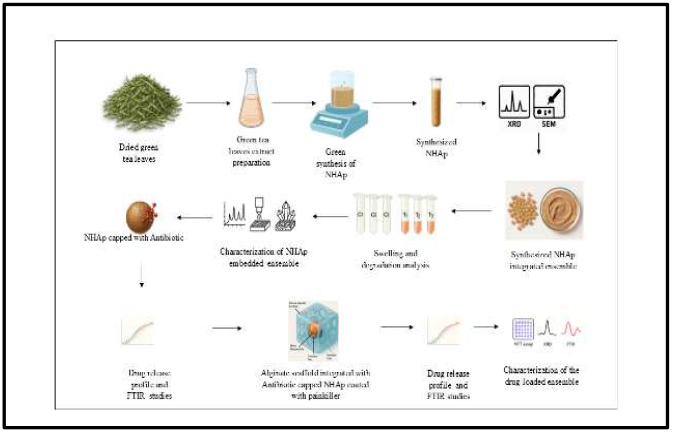

## 1) Introduction

Dental injuries are a global health concern impacting almost a billion people worldwide. Timely and appropriate treatment is crucial to treat dental injuries, along with preventive measures such as mouthguards during sports are crucial. Understanding clinical patterns and epidemiology is crucial for developing targeted management strategies, including advances in dental tissue engineering**[1,4]**.

Dental bone tissue engineering, a domain in regenerative dentistry, focuses on the repair or regeneration of damaged or lost dental bone. It combines materials science, biology, and engineering to innovate scaffolds, growth factors, and cell-based therapies that restore structure and function **[5]**. This technique promotes biological integration, nerve regeneration, and vascularization, along with minimally invasive and customized solutions catering to patient-specific needs **[6]**.

Polymers play a key role because of their tunable chemistry and structural properties **[7]**. Natural and synthetic polymers, as well as composites, impact biocompatibility, degradation, and scaffold design **[8,12]**. Alginate favours regeneration of alveolar bone through osteogenic and customizable properties **[13]**.

Hydroxyapatite mimics the natural bone mineral, enhancing biocompatibility and osteogenesis **[14]**. Phytochemical-mediated synthesis, specifically utilizing green tea extract as a capping agent in the synthesis of nHAp, enhances its osteogenic property, biocompatibility, its chemical stability, contributes towards its antimicrobial property, and makes this synthesis approach eco-friendly, making it highly suitable for dental applications **[15,16]**.

## 2) Materials and methods

### 2.1) Materials

Ammonium dihydrogen orthophosphate was procured from Karnataka Fine Chem, Sodium alginate was procured from SRL chemicals Ltd, Calcium chloride was procured from SDFCL Ltd, Mumbai. Tetley Green tea long leaf original was used for the extract preparation.

### 2.1) Nanohydroxyapatite (nHAp) synthesis

#### Extract preparation

Green tea Leaves were finely ground into powder form using a mixer.10 mg of this leaf powder was added to 50 ml of deionized water & boiled for 2 hours. Once cooled, the mixture was filtered using Wattmann filter paper, and the extract obtained was stored at 4^°^C for further experiments.

#### Green synthesis of nHAp

10 ml of 0.1M Calcium chloride was added to 10 ml of 0.6M Ammonium dihydrogen orthophosphate solution and mixed using a magnetic stirrer to form a homogenous mixture. This was followed up by making the pH to 12 using 0.8 M NaOH, after which the mixture was added with 5 ml of the green tea leaf extract and was mixed using the magnetic stirrer for 1hour and left undisturbed at room temperature to get gelatinous precipitate. The obtained gelatinous ppt was dried in a microwave oven for 10 minutes and calcinated in a 100^°^C furnace for 1 hour. The dried cake was crushed to derive nanohydroxyapatite **[17]**.

#### Characterization of nHAp

The synthesised nHAp was characterised by X-ray diffraction to confirm its molecular composition. The nHAp was tested at room temperature using a Panalytical diffractometer (Xpert Pro powder). (Cu K_ radiations) working at a voltage of 40 kV. XRD was carried out at a 2θ angle range of 10–70^°^, and the process parameters were kept as: scan step time 0.05 s and scan step size 0.02.

### 2.2) Preparation of scaffold integrated with nHAp

#### Optimization of the alginate concentrations

4%, 6%, and 8% sodium alginate solutions were prepared. They were individually mixed with 10 mg of nanohydroxyapatite and poured into separate Petri plates. They were polymerised by adding CaCl_2_ solution dropwise with constant stirring to ensure uniform solidification. The samples were kept in the freezer for 24 hours and then subjected to lyophilisation.

#### Preparation of nHAp-embedded alginate scaffold

After optimisation, 8% alginate was chosen for further experiments as it showed the most promising mechanical properties. For scaffold preparation, a mixture of 8% alginate solution was used, and 10 mg of nHAp was added to it and subjected to continuous stirring using a magnetic stirrer. This mixture was added with chilled CaCl_2_ for crosslinking **[18]**. After 15 minutes of incubation at room temperature, the sample was lyophilised to obtain scaffolds of the same.

### 2.3) Characterization of Scaffold

#### XRD analysis

The scaffolds fabricated were subjected to XRD analysis. The structure of the composite scaffold was analysed with the help of XRD studies. XRD patterns of the composite scaffolds were evaluated at standard conditions using a Panalytical diffractometer (XPERT Pro powder). (Cu K radiations) operating at a voltage of 40 kV. XRD was carried out for the 2-theta angle range of 10– 70^°^, and the process parameters were kept as follows: 0.05 s for the scan step time and 0.02 for the scan step size.

#### Swelling analysis

The 8% alginate scaffold without the integration of nHAp was taken as the control sample, and the scaffold with the integration of nHAp was taken as the Test sample. The dry weights of the scaffold were noted. Scaffolds are then placed into the prepared PBS solution separately. The wet weight scaffolds were weighed for a week (Ex: Day 1,3,5, and 7) and a graph of weight v/s day was plotted **[19]**. The swelling percentage was calculated using equation 1.

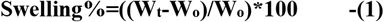

Wherein W_t_ is the wet weight of the scaffold, W_o_ is the initial weight of the scaffold.

#### Degradation studies

The 8% alginate scaffold without the integration of nHAp was taken as the control sample, and the scaffold with the integration of nHAp was taken as the Test sample. The dry weights of the scaffold were taken before the experiment. The pre-weighed scaffolds were added to the PBS buffer solution with lysozyme. The scaffolds were weighed at Days 5,7,9,11, and 15. A weight v/s day graph is plotted for the degradation ratio values **[20]**. The degradation ratio was calculated using equation 2.

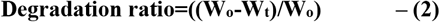

Wherein W_t_ is the wet weight of the swollen scaffold, W_o_ is the initial weight of the dry scaffold.

### 2.4) Drug loading and release studies

#### Preparation of a standard curve for antibiotic and painkiller

Standard graphs of the chosen antibiotic (Amoxycillin) and painkiller (Paracetamol) were individually plotted for concentrations 0.25,0.5,1,1.5 mg/ml at 270 and 196 nm, respectively.

#### Capping of Antibiotic to nHAp

An optimised concentration of Amoxicillin solution was prepared and used for capping Amoxicillin onto nHAp in ratios 1:1 and 1:2. The amoxicillin-capped nHAp was dried and integrated into the alginate scaffolds individually and subjected to lyophilization.

#### Antibiotic (Amoxicillin) release profile studies

For evaluating the release profile of antibiotics, the antibiotic-capped nHAp was immersed in PBS solution, and spectrometric analysis of the solution was carried out at 270 nm for time intervals of 15 minutes till 150 minutes to study the Amoxicillin release profile [**21]**

#### Preparation of scaffold encapsulated with antibiotic-capped nHAp and painkiller

20 ml of 8% sodium alginate solution was taken, and 10 mg of the prepared Amoxicillin-capped nHAp was added and mixed thoroughly. The mixture was added dropwise into chilled calcium chloride, and then lyophilised (as per the protocol given above) to prepare the scaffolds. Thus prepared scaffolds were incubated for 2 hours in10ml of painkiller (Paracetamol) solution at a concentration of 1mg/ml. After incubation, the scaffolds were removed from the painkiller solution and air-dried until they were dry.

#### Paracetamol release profile studies

The scaffolds coated with paracetamol were immersed in PBS solution, and the Paracetamol release into PBS was noted at 196 nm **[22]**.

### 2.5) Biomineralization studies

A 100 mL solution of simulated body fluid (SBF) was prepared using the following reagents: 0.745 g NaCl, 0.044 g KCl, 0.035 g NaHCO3, 0.116 g CaCl2·2H2O, 0.031 g MgSO4·7H2O, 0.023 g Na2HPO4·2H2O, and 0.15 g Tris base. These reagents were dissolved in 90 mL of deionized water under constant magnetic stirring. The pH of the solution was adjusted to 7.4 using 1 M HCl. Subsequently, the solution was brought to a final volume of 100 mL with deionized water. Scaffolds embedded with Green tea nHAp were immersed in the prepared SBF solution. Following a designated incubation period (14 days), the SBF was aspirated, and the scaffolds were lyophilized and subjected to SEM analysis. The SEM analysis was carried out using Ultra55 FE-SEM Carl Zeiss EDS **[23]**.

### 2.6) Cytocompatibility assay

The MTT analysis was carried out to evaluate the cytocompatibility of the cells after interacting with the scaffold samples:

- nHAp + Alginate scaffold.
- nHAp + Alginate + Antibiotic scaffold.
- nHAp+ Alginate + Antibiotic + Painkiller scaffold

The MTT assay determines the extent to which viable cells can cause the conversion of tetrazolium in MTT to formazan. The assay was done on L929(Mouse fibroblast cell lines), MG 63 (Osteosarcoma cell lines), and POB (Primary Osteoblast cells) were cultured under standard culturing conditions.

Following a seeding density of 104 cells per well into 96-well plates, under sterile circumstances, the scaffold with nHAp, antibiotics, and painkillers in different permutations and combinations was kept immersed in full medium at 37 °C and 48 hours with agitation. The medium consisting of the leachable was gathered in a Falcon tube following the incubation period **[24]**.

### 2.7) FTIR analysis

The FTIR analysis was carried out for the nHAp embedded alginate scaffold and the nHAp embedded alginate scaffold integrated with paracetamol and antibiotic. FTIR was carried out using a Perkin Elmer Frontier FTIR spectrophotometer operating in the wavelength range 4000-600 cm^-1^ to study whether the chemical integrity of the nHAp alginate scaffold was interfered with by the drugs integrated with the scaffold. The method followed was ATR method **[25]**.

### 2.8) ALP analysis

The alkaline phosphatase (ALP) activity assay was carried out to understand the biological effects of a scaffold. Cells, including odontoblast-like cells (OLC), primary osteoblasts (POB), and their co-culture (OLC+POB), were seeded onto pre-sterilized scaffolds and maintained under standard culture conditions in sustained growth media. At specific time intervals (day 7, day 14, and day 21), cells were subjected to enzymatic detachment from the scaffolds using 0.25% trypsin-EDTA solution to ensure efficient recovery of adherent cells. The collected cell suspensions were subjected to DNA isolation using a standard genomic DNA extraction protocol to obtain high-quality nucleic acids. The extracted DNA was quantified spectrophotometrically to determine concentration and purity, which served as an indirect measure of cellular proliferation and viability at each time point **[26]**.

### 2.9) Statistical Analysis

Experiments were carried out in triplicate, for which standard errors were calculated, and graphs were plotted.

## 3) Results and Discussion

### 3.1) Nanoparticle synthesis: Green synthesis of nanohydroxyapatite(nHAp)

nHAp was synthesized using Green tea leaf extracts by the green synthesis approach as stated in the above methodology (**Figure 1**).

**Figure 1:**
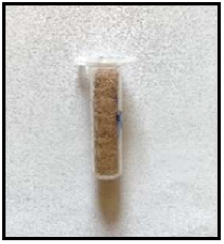
nHAp synthesized using Green tea leaf extract.

#### Characterization of synthesized nHAp

XRD of nHAp showed sharp peaks at crystal planes (002),(112), at 2θ = 26.0° and 32.1° respectively, matching the Joint Committee on Powder Diffraction Standards (JCPDS) standard PDF card No. 01-86-1199, confirming the elemental composition for hydroxyapatite (**Figure 2**).

**Figure 2:**
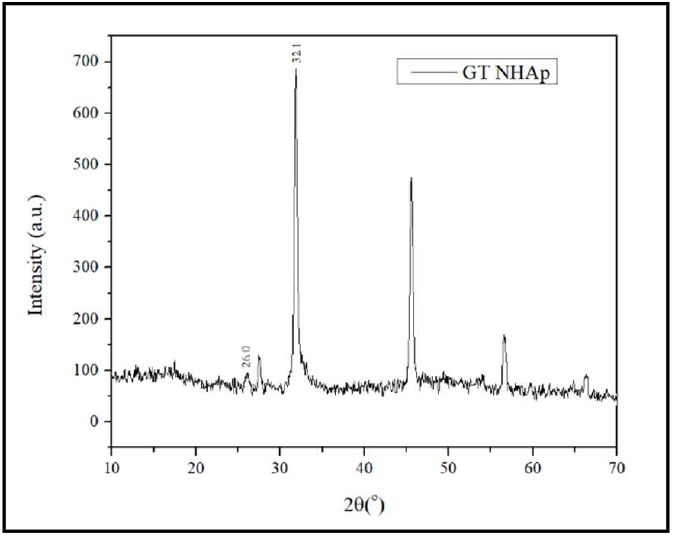
XRD results of nHAp synthesized using Green tea leaves extract.

### 3.2) Preparation of scaffold integrated with nanohydroxyapatite

The scaffold (**Figure 3**) was prepared according to the protocol stated above.

**Figure 3:**
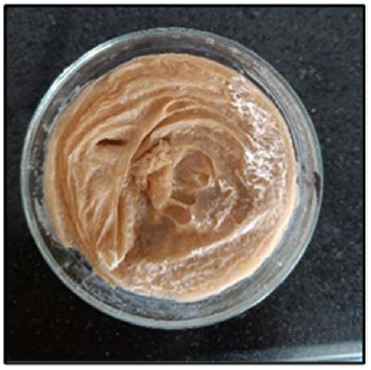
nHAp embedded alginate scaffold.

#### Optimization of the alginate concentrations

The alginate concentration optimisation studies were carried out, as it is important to tailor the biological properties and physicochemical properties to application-specific requirements. Alginate’s concentration impacts swelling behavior, porosity, mechanical strength, and drug release properties, which directly influence the scaffold’s efficiency and efficacy in tissue engineering. The alginate scaffolds of 4%,6%, and 10% were integrated with nHAp and prepared as per the above protocol. The wet weights of the scaffolds were taken on Days 1,3,5,7 to check for their swelling capabilities. The optimisation studies revealed that 8% alginate is ideal for the preparation of scaffolds for further studies (**Figure 4**)

**Figure 4:**
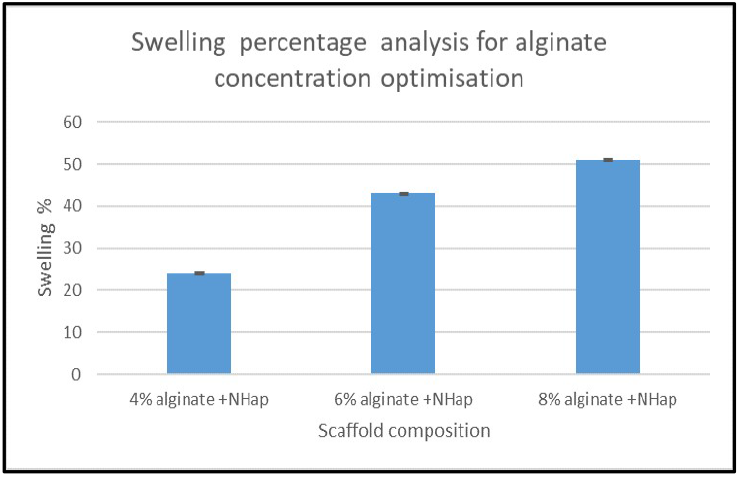
Swelling ratio for different concentrations on day 7.

### 3.3) Characterization of Scaffold using XRD

The XRD of pure alginate shows a broad, Diffuse peak instead of a well-defined, sharp peak. This characteristic is mainly due to the amorphous nature of the alginate **(Figure 5)**. Such a structure is typical for natural polysaccharides like alginate, coinciding with the previously reported results for Sodium Alginate, which show a broad hump in the 2θ range of approximately 10-30^°^ °. This amorphous behavior is advantageous for tissue engineering applications, as it often leads to flexibility and high porosity **[27]**.

**Figure 5:**
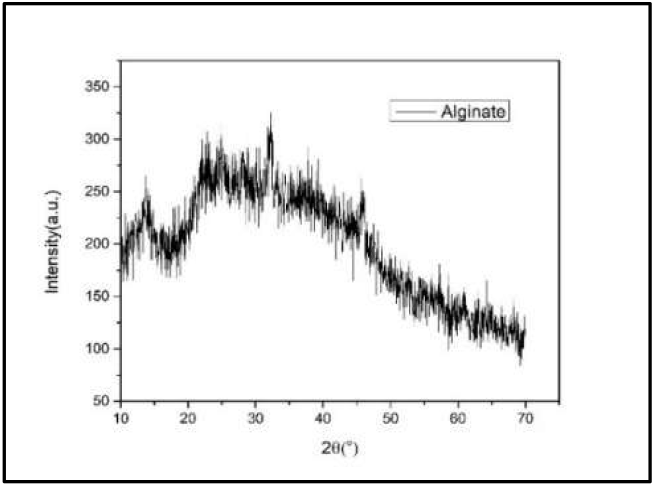
XRD analysis of alginate.

In contrast, the XRD results of the composite scaffold reveal sharp peaks at 32^°^, 45^°^, and 56^°^ that correlate with the peaks of Crystalline nanohydroxyapatite. The retention of the broad background peaks further shows that the composite contains both alginate and crystalline HAp.This indicates successful integration of the nHAp into the alginate scaffold. This Combination of amorphous and crystalline nature demonstrates that the composite has the advantages of porosity and flexibility of the alginate along with the bioactivity and structural integrity of the hydroxyapatite (**Figure 6**) [**28**].

**Figure 6:**
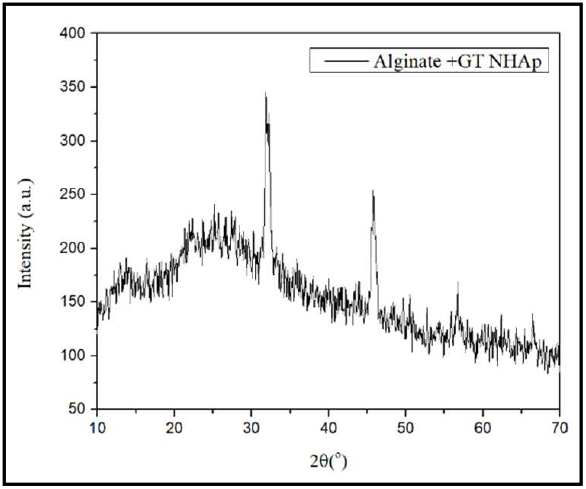
XRD of alginate +Green Tea NHAp.

#### SEM analysis

The SEM micrograph at 10.00 KX magnification shows successful integration of Green tea-mediated nHAp within the alginate scaffold, exhibiting uniform distribution, almost spherical nanoparticles with low agglomeration. The porous microstructure depicts effective scaffold formation, which can be beneficial for dental bone tissue engineering by enhancing cell adhesion and nutrient diffusion. The enhanced morphological characteristics associated with green tea derivatives indicate enhanced bioactivity and Osteoconductivity, supporting the composite’s applicability in dental bone tissue engineering **[29]**(**Figure 7**).

**Figure 7:**
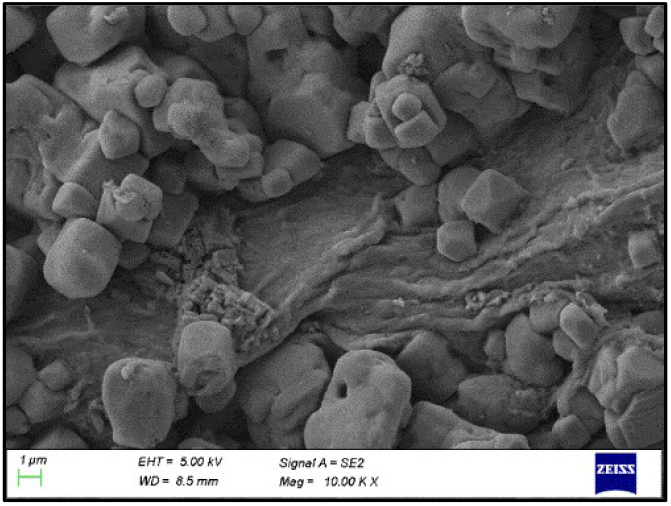
SEM image of green tea nHAp embedded in polymeric scaffold.

#### Swelling analysis

Swelling analysis was carried out to evaluate the fluid uptake ability and understand the mechanical properties of the scaffold when alginate was integrated with the nanohydroxyapatite in comparison to just the alginate to validate the impact of the integration of nHAp on influencing the scaffold’s performance. The Swelling studies were carried out as explained earlier, and it was observed that the scaffolds integrated with the nHAp synthesized using green tea extract gave 12 % better swelling by day 7 in comparison to the control scaffold, showing only 12%, indicating that the integration of nHAp improved the swelling property (**Figure 8**).

**Figure 8:**
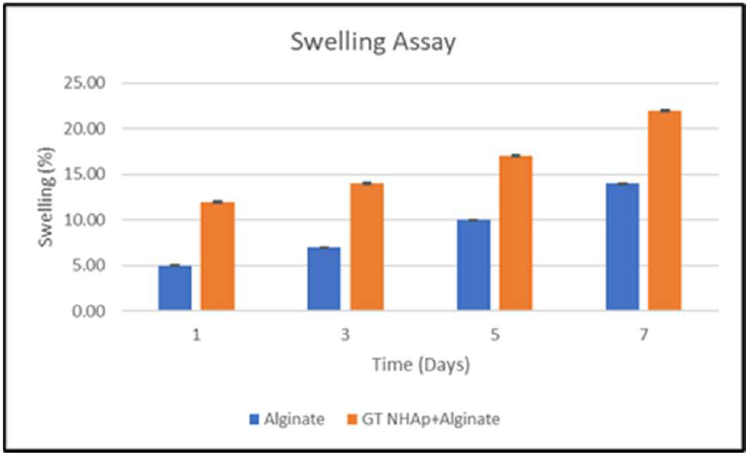
Swelling Assay results.

##### 3.2.5) Degradation Analysis

Degradation of the scaffolds is a critical criterion for tissue engineering design. Preferably, the scaffold’s degradation rate should be similar to the rate at which the new tissue forms to ensure proper functionalization of the newly formed tissue. Upon taking wet weights on Days 5, 7, 9, 11, and 15, it was observed that the scaffold weight increased for the first 24 hours for both the control (alginate) and the test (alginate + nHAp), and later started degrading. After day 9, it was found that the degradation was accelerated in the control, while the degradation in the test sample was happening at a controlled and slower pace. Upon calculating the degradation ratio at day 15, it was found that the control had a higher degradation ratio compared to the test, indicating that the addition of nHAp slowed down the degradation process of the scaffold. From this assay, we can confirm that the addition of nHAp extends the time of degradation of the scaffold synthesized (**Figure 9**).

**Figure 9:**
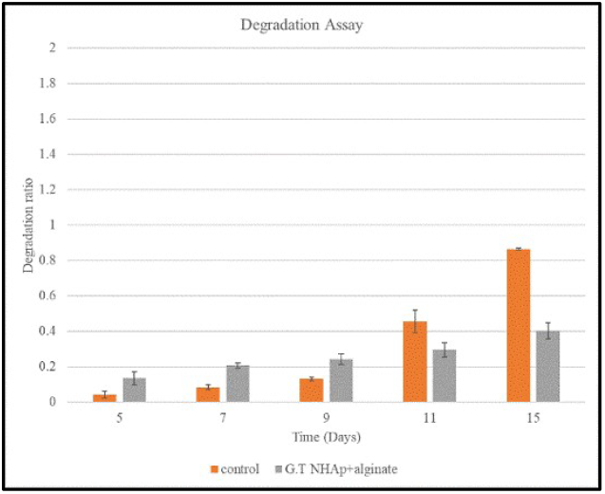
Graph for degradation analysis.

### 3.4) Preparation of standard curve for antibiotic and painkiller

#### Antibiotic

For the drug loading studies, the antibiotic drug Amoxicillin was chosen, due to its extensive use as a dental antibiotic along with its solubility in organic solvents and distilled water, which was found to be 1mg/ml. This value was obtained after a standard graph **Figure 10**. OD v/s concentration of the drug was carried out for concentrations 0.5mg/ml to 2mg/ml at an absorbance reading of 270nm.

**Figure 10:**
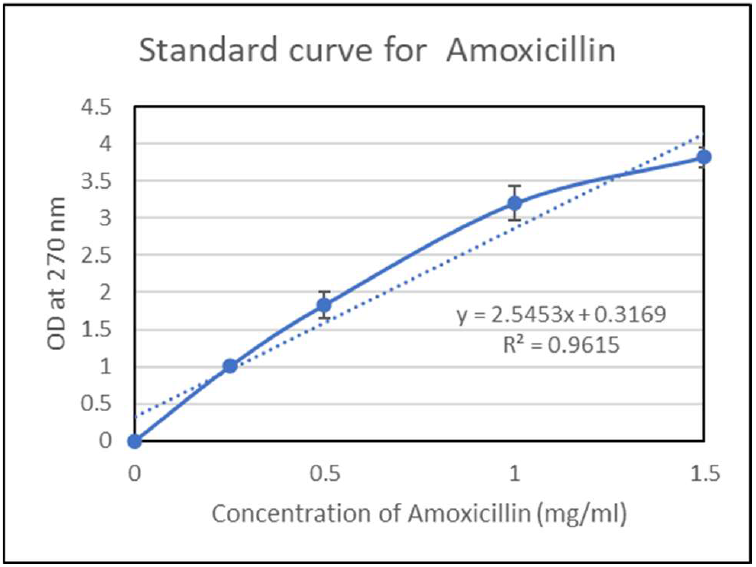
Standard Absorption Curve for Amoxicillin at O.D. of 270nm.

#### Painkiller

For the drug loading, the painkiller chosen was Paracetamol, as it is one of the most commonly found and used painkillers in the market. It also has a high solubility in organic solvents as well as in distilled water. The solubility of paracetamol in distilled water was determined to be 1 mg/ml. This was established by measuring the absorbance of various concentrations of paracetamol solutions in the intervals between 0.25mg/ml to 1 mg/ml **(Figure 11)**.

**Figure 11:**
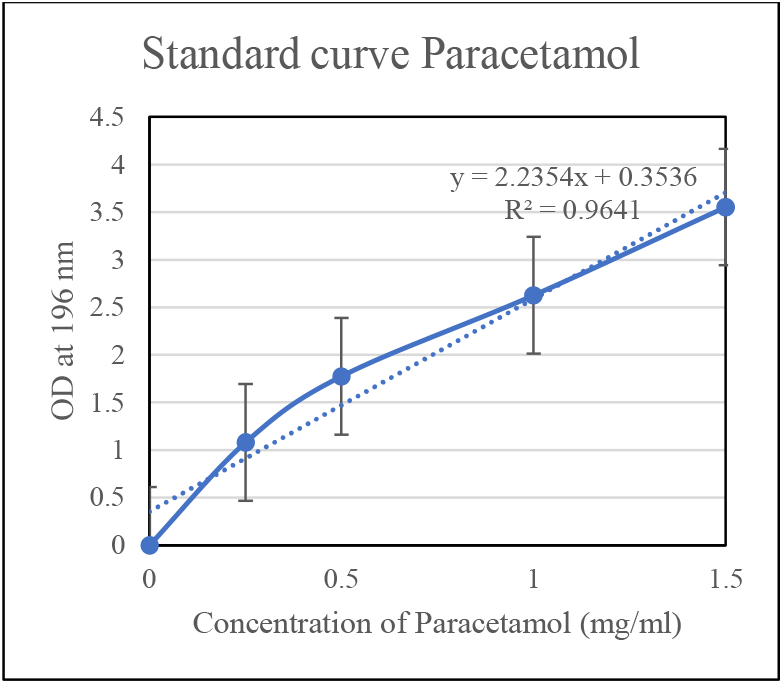
Standard Absorption Curve for Paracetamol at O.D. of 196nm.

### 3.4) Drug Loading and Release Profile

#### Antibiotic release profile

The Amoxicillin release was found to be sustained in both scaffolds. Notably, the scaffold with a 1:2 ratio of Amoxicillin to nHAp showed a higher cumulative release in comparison to the scaffold with a 1:1 ratio of Amoxicillin to nHAp, which might be due to increased nanohydroxyapatite content, which in turn provides greater drug loading capacity. The results indicate that adjusting the antibiotic to nanohydroxyapatite ratio is an effective strategy to optimise the release rate and achieve higher antibiotic availability in a dual drug delivery system (**Figure 12**).

**Figure 12:**
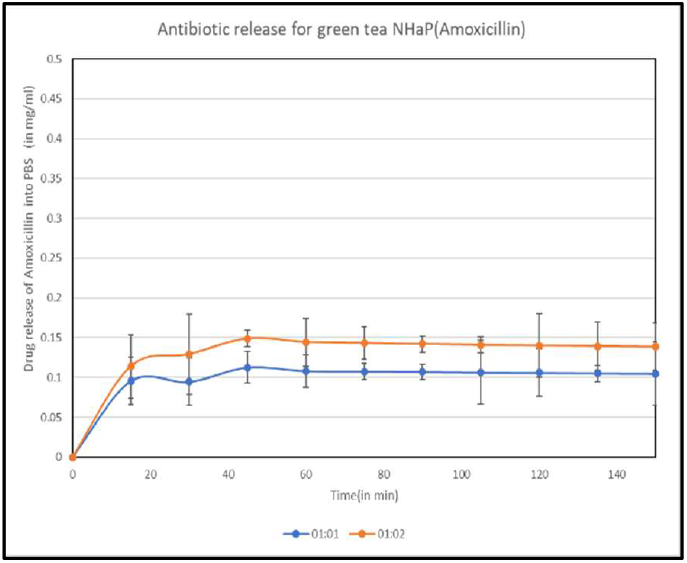
Graph for antibiotic release profile of Antibiotic-capped Green tea nHAp.

#### Painkiller release profile

From the painkiller (paracetamol) release profile scaffold, we can infer that there was a rapid release of the painkiller for the first 15 minutes, followed by a slow and steady decline that corresponds to the burst release profile of the painkiller from the drug-loaded scaffold (**Figure 13**).

**Figure 13:**
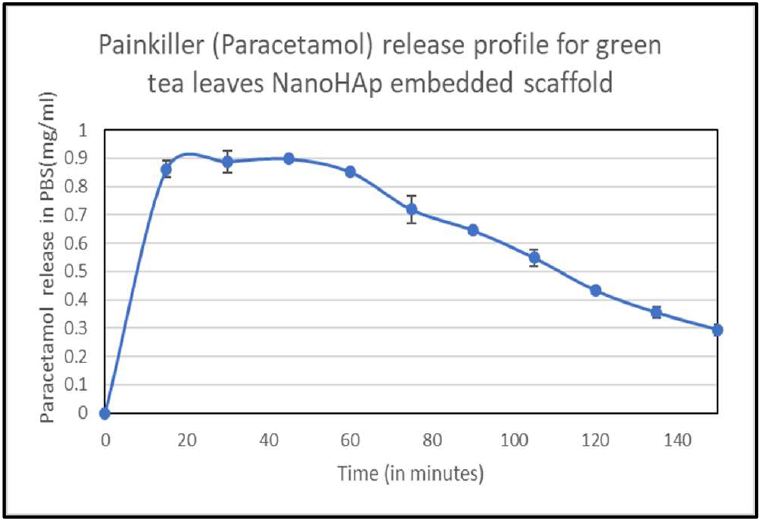
Painkiller release profile for green tea leaves nHAp embedded scaffold.

### 3.5) Biomineralization studies

Biomineralization studies were carried out to evaluate the scaffold’s capacity to induce the formation of bone-like apatite on its surface, which validates that the scaffold is suitable for bone regeneration applications. The SEM image (Figure 15) of green tea nHAp-embedded alginate scaffold, following 14 days of incubation in simulated body fluid (SBF), shows a densely mineralized surface characterized by uniform deposition of spherical apatite particles. This appreciable level of apatite formation depicts the scaffold’s enhanced bioactivity and ability to support mineral nucleation, highlighting its suitability for dental bone tissue engineering applications. The observed morphology aligns with established evidence that SBF-immersion enhanced calcium-phosphate layer development, thereby promoting Osteoconductivity and osteointegration potential (**Figure 14**).

**Figure 14:**
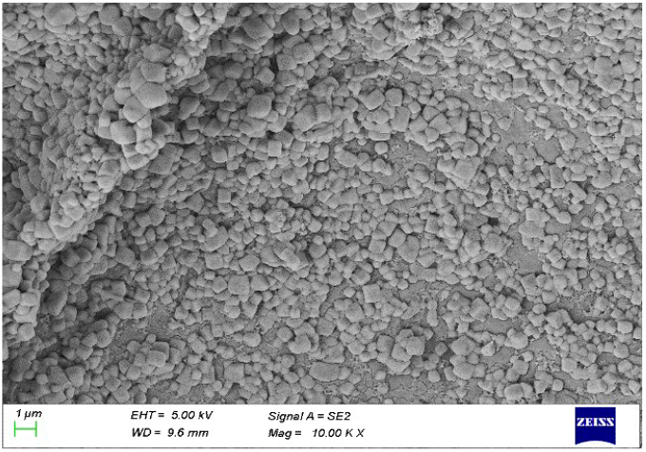
SEM image of the green tea nHAp embedded alginate scaffold immersed in the SBF solution for 14 days.

**Figure 15:**
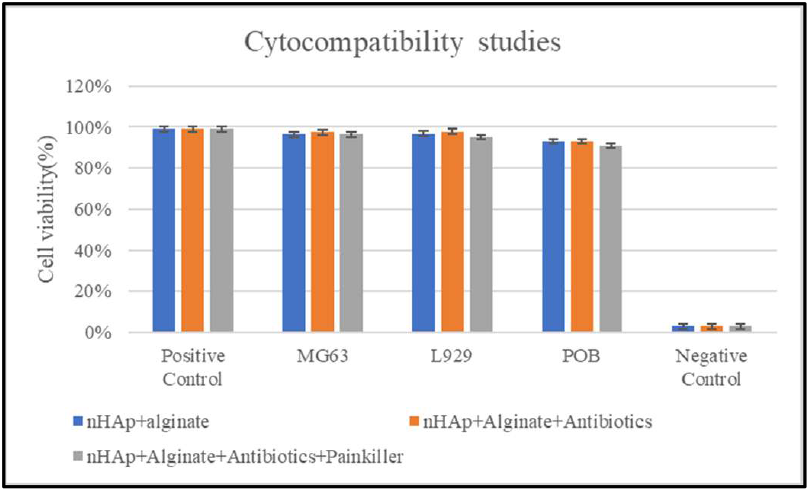
Cell Viability of the samples in different cell lines.

### 3.6) Cytocompatibility assay

MTT assay was used to evaluate the cytocompatibility of the nHAp coated with antibiotics and encapsulated inside an alginate sphere. This whole ensemble was coated with painkillers in this study.

The findings showed that post-48-hour incubation, 93% of the different osteoblastic cell lines were compatible with the nanocomposite scaffold. Comparing the alginate scaffolds used as a reference to the alginate/nHAp composite scaffolds, this result implies that there is not much of a hazardous leachable. These findings demonstrate that the Green tea nHAp-alginate scaffold is biocompatible with all varieties of osteoblastic lineage cells (**Figure 15**).

### 3.7) FTIR Analysis

FTIR was carried out to understand if the integration of drugs interferes with the chemical integrity of the scaffold. FTIR analysis of the NanoHAp alginate scaffold and drug-loaded composites validated the successful integration of both scaffold components and drugs without compromising chemical integrity. Characteristic peaks at 3372 and 3351 cm^−1^ correspond to O– H/N–H stretching vibrations from alginate hydroxyl groups and structural water in nanohydroxyapatite. Specific peaks at 1633 and 1628 cm^−1^ represent C=O stretching of amide I and carboxylate groups in alginate, while phosphate peaks specific to nHAp appear near 1027–1028 cm^−1^ and 669–666 cm^−1^. The integration of paracetamol and amoxicillin introduced additional bands at 1511 and 1410 cm^−1^, typically aromatic C=C stretching and amide II vibrations from the drugs. Amoxicillin further contributes O–H and N– H stretches within 3200–3500 cm^−1^, β-lactam and amide carbonyl stretches between 1800–1650 cm^−1^, and N–H bending and C–N stretching in the range 1300–1250 cm^−1^, while paracetamol shows amide C=O and aromatic vibrations consistent with its structure. The absence of peak loss or new covalent bond formation in the drug-loaded scaffold indicates physical entrapment and hydrogen bonding rather than chemical alteration, thereby validating the scaffold’s structural stability and its applicability in dental bone tissue engineering and for controlled drug delivery **(Figure 16)[30,32]**.

**Figure 16:**
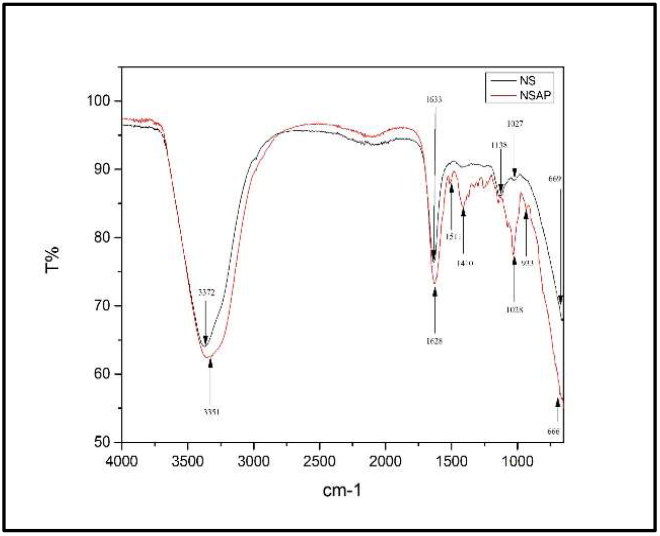
FTIR analysis of nHAp embedded alginate scaffold(NS) and nHAp embedded alginate scaffold integrated with Amoxicillin and Paracetamol(NSAP).

### 3.8) ALP Activity

The ALP activity graph projects a time-dependent enhancement in osteogenic differentiation across all tested groups, with the co-cultured OLC+POB group consistently projected to have higher ALP activity in comparison to the single cell lines (OLC and POB) at each time point. On day 7, ALP levels are minimal for all groups, but by day 14 and day 21, a noticeable increase is observed, particularly for the co-culture, which achieves the greatest activity by day 21. This trend indicates that the scaffold leachate treatment effectively promotes osteoblastic activity and matrix maturation, with synergistic effects evident in the mixed cell population. These inferences suggest that co-culturing OLC and POB enhances osteogenic response—likely due to cell-cell interactions—and highlights the potential of the scaffold as a potent inducer of bone tissue formation. The results align with the expected biological pattern of ALP activity as an early marker for osteoblast differentiation and show the added value of using co-culture systems for evaluating novel biomaterials in regenerative applications **(Figure 17)**.

**Figure 17:**
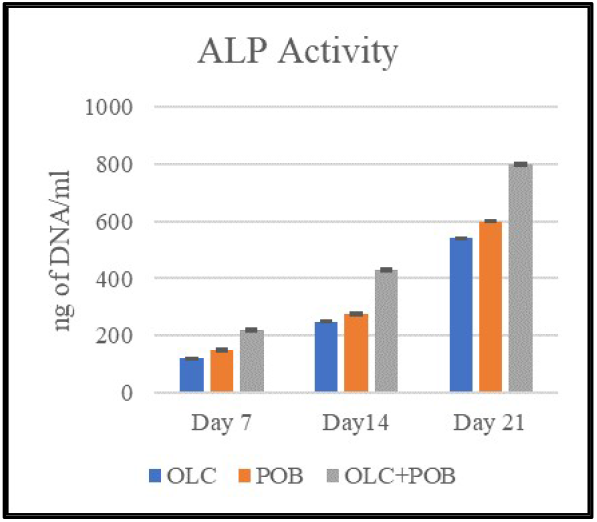
ALP activity assay for nHAp embedded scaffold.

## 4) CONCLUSION

Traumatic dental tissue, being a global health concern, impacts a spectrum of age groups globally, highlighting the significance of personalised therapies to treat the same. Dental tissue engineering offers personalized treatments that cater to patient-specific needs.

This research is focused on developing a nanocomposite scaffold for dental bone tissue engineering with dual drug delivery functionality. The scaffold is designed for the rapid release of painkiller and sustained release of antibiotic to deal with inflammation and infection. Integration of hydroxyapatite enhances osteogenesis, which promotes bone regeneration. Nanohydroxyapatite was synthesized using the green synthesis process and was confirmed by XRD analysis. Swelling and degradation analysis showed that the integration of nanohydroxyapatite enhanced the swelling properties and decreased the rate of degradation, supporting bone healing over a longer time duration.

Drug release profiles showed that antibiotic-capped nanohydroxyapatite provides controlled release, with encapsulation ratio adjustments as a tool for optimizing the treatments. This ensures the antibiotic release is precise and sustained, while the rapid release of the painkiller deals with inflammation.MTT assay performed validated the cytocompatibility, showing 93% compatibility for osteoblast and fibroblast cell lines. ALP activity studies confirmed the osteogenic potential of the scaffold, hence validating its use in bone regeneration applications. This system can be tested with different drugs for various dental applications and could be integrated with osteoblastic stem cells. Further safety testing in animal models can also be carried out in mouse models.

## CRediT authorship contribution statement

Ananya N. Nayak: Conceptualization, Methodology, Investigation, Supervision, Formal analysis, Data curation, Visualization, Writing – original draft. Roshni Ramachandran: Conceptualization, Validation, Supervision, Visualization, Writing – review & editing, Interpretation of results. Hansel P. Shah: Investigation, Data curation. Ishani Chowdhary: Investigation, Data curation. Punya Sharma: Investigation, Data curation.

## References

[1] L.V. Reddy, R. Bhattacharjee, E. Misch, M. Sokoya, Y. Ducic, Dental injuries and management, Facial Plast. Surg. 35(6) (2019) 607–613. 10.1055/s-0039-1700877.

[2] S. Petti, J.O. Andreasen, U. Glendor, L. Andersson, NA0D – the new traumatic dental injury classification of the World Health Organization, Dent. Traumatol. 38(3) (2022) 170– 174. 10.1111/edt.12753.

[3] L. Mordini, P. Lee, R. Lazaro, R. Biagi, L. Giannetti, Sport and dental traumatology: surgical solutions and prevention, Dent. J. 9(3) (2021) 33. 10.3390/dj9030033.

[4] C. Bourguignon, N. Cohenca, E. Lauridsen, M.T. Flores, A.C. O’Connell, P.F. Day, … L. Levin, International Association of Dental Traumatology guidelines for the management of traumatic dental injuries: 1. Fractures and luxations, Dent. Traumatol. 36(4) (2020) 314–330. 10.1111/edt.12578.

[5] S. Joshi, S. Eshwar, V. Jain, Marine polysaccharides: biomedical and tissue engineering applications, in: A. Choi, B. Ben-Nissan (Eds.), Marine-Derived Biomaterials for Tissue Engineering Applications, Springer Ser. Biomater. Sci. Eng., Vol. 14, Springer, Singapore, 2019, pp. 481–502. 10.1007/978-981-13-8855-2_19.

[6] M. Saleh Hasani Jebelli, A. Yari, N. Nikparto, S. Cheperli, A. Asadi, A.A. Darehdor, … L.K. Hakim, Tissue engineering innovations to enhance osseointegration in immediate dental implant loading: a narrative review, Cell Biochem. Funct. 42(2) (2024) e3974. 10.1002/cbf.3974.

[7] Z. Terzopoulou, A. Zamboulis, I. Koumentakou, G. Michailidou, M.J. Noordam, D.N. Bikiaris, Biocompatible synthetic polymers for tissue engineering purposes, Biomacromolecules 23(5) (2022) 1841–1863. 10.1021/acs.biomac.2c00047.

[8] D.T. Wu, J.G. Munguia-Lopez, Y.W. Cho, X. Ma, V. Song, Z. Zhu, S.D. Tran, Polymeric scaffolds for dental, oral, and craniofacial regenerative medicine, Molecules 26(22) (2021) 7043. 10.3390/molecules26227043.

[9] D. Yang, J. Xiao, B. Wang, L. Li, X. Kong, J. Liao, The immune reaction and degradation fate of scaffold in cartilage/bone tissue engineering, Mater. Sci. Eng. C 104 (2019) 109927. 10.1016/j.msec.2019.109927.

[10] F. Zhang, M.W. King, Biodegradable polymers as the pivotal player in the design of tissue engineering scaffolds, Adv. Healthc. Mater. 9(13) (2020) 1901358. 10.1002/adhm.201901358.

[11] B. Özcolak, B. Erenay, S. Odabaş, K.D. Jandt, B. Garipcan, Effects of bone surface topography and chemistry on macrophage polarization, Sci. Rep. 14(1) (2024) 12721. 10.1038/s41598-024-62484-3.

[12] T. Chen, Y. Jiang, J.P. Huang, J. Wang, Z.K. Wang, P.H. Ding, Essential elements for spatiotemporal delivery of growth factors within bio-scaffolds: a comprehensive strategy for enhanced tissue regeneration, J. Control. Release 368 (2024) 97–114. 10.1016/j.jconrel.2024.02.006.

[13] D.R. Sahoo, T. Biswal, Alginate and its application to tissue engineering, SN Appl. Sci. 3(1) (2021) 30. 10.1007/s42452-02004096-w.

[14] S.U. Rahman, Hydroxyapatite and tissue engineering, in: Handbook of Ionic-Substituted Hydroxyapatites, Woodhead Publ., 2020, pp. 383– 400. 10.1016/B978-0-08-102834-6.00016-1.

[15] S. Singh, A. Pal, S. Mohanty, Nano structure of hydroxyapatite and its modern approach in pharmaceutical science, Res. J. Pharm. Technol. 12(3) (2019) 1463–1472. 10.5958/0974-360X.2019.00243.9.

[16] A. Siriphap, A. Kiddee, A. Duangjai, A. Yosboonruang, G. Pook-In, S. Saokaew, … A. Rawangkan, Antimicrobial activity of the green tea polyphenol (–)-epigallocatechin-3-gallate (EGCG) against clinical isolates of multidrug-resistant Vibrio cholerae, Antibiotics 11(4) (2022) 518. 10.3390/antibiotics11040518.

[17] D. Govindaraj, M. Rajan, Synthesis and spectral characterization of novel nanohydroxyapatite from Moringa oleifera leaves, Mater. Today Proc. 3(6) (2016) 2394–2398. 10.1016/j.matpr.2016.04.153.

[18] V. Jayachandran, S.S. Murugan, P.A. Dalavi, Y.D. Vishalakshi, G.H. Seong, Alginate-based composite microspheres: preparations and applications for bone tissue engineering, Curr. Pharm. Des. 28(13) (2022) 1067–1081. 10.2174/1381612828666220518142911.

[19] O.D. Frenţ, N. Duteanu, A.C. Teusdea, S. Ciocan, L. Vicaş, T. Jurca, … E. Marian, Preparation and characterization of chitosan– alginate microspheres loaded with quercetin, Polymers 14(3) (2022) 490. 10.3390/polym14030490.

[20] J. Kurowiak, A. Kaczmarek-Pawelska, A.G. Mackiewicz, R. Bedziński, Analysis of the degradation process of alginate-based hydrogels in artificial urine for use as a bioresorbable material in the treatment of urethral injuries, Processes 8(3) (2020) 304. 10.3390/pr8030304.

[21] X. Mo, D. Zhang, K. Liu, X. Zhao, X. Li, W. Wang, Nano-hydroxyapatite composite scaffolds loaded with bioactive factors and drugs for bone tissue engineering, Int. J. Mol. Sci. 24(2) (2023) 1291. 10.3390/ijms24021291.

[22] N.T.T. Uyen, Z.A.A. Hamid, N.B. Ahmad, Synthesis and characterization of curcumin-loaded alginate microspheres for drug delivery, J. Drug Deliv. Sci. Technol. 58 (2020) 101796. 10.1016/j.jddst.2020.101796.

[23] D. Bellucci, R. Salvatori, A. Anesi, L. Chiarini, V. Cannillo, SBF assays, direct and indirect cell culture tests to evaluate the biological performance of bioglasses and bioglass-based composites: three paradigmatic cases, Mater. Sci. Eng. C 96 (2019) 757–764. 10.1016/j.msec.2018.12.006.

[24] Q. Zhou, T. Wang, C. Wang, Z. Wang, Y. Yang, P. Li, … L. Nie, Synthesis and characterization of silver nanoparticles-doped hydroxyapatite/alginate microparticles with promising cytocompatibility and antibacterial properties, Colloids Surf. A Physicochem. Eng. Asp. 585 (2020) 124081. 10.1016/j.colsurfa.2019.124081.

[25] F. Zapata, A. Lopez-Fernandez, F. Ortega-Ojeda, G. Quintanilla, C. Garcia-Ruiz, G. Montalvo, Introducing ATR-FTIR spectroscopy through analysis of acetaminophen drugs: practical lessons for interdisciplinary and progressive learning for undergraduate students, J. Chem. Educ. 98(8) (2021) 2675–2686. 10.1021/acs.jchemed.0c01231.

[26] M.K. Haider, D. Kharaghani, L. Sun, S. Ullah, M.N. Sarwar, A. Ullah, … I.S. Kim, Synthesized bioactive lignin nanoparticles/polycaprolactone nanofibers: a novel nanobiocomposite for bone tissue engineering, Biomater. Adv. 144 (2023) 213203. 10.1016/j.bioadv.2022.213203.

[27] C. Larosa, M. Salerno, J.S. de Lima, R.M. Meri, M.F. da Silva, L.B. de Carvalho, A. Converti, Characterisation of bare and tannase-loaded calcium alginate beads by microscopic, thermogravimetric, FTIR and XRD analyses, Int. J. Biol. Macromol. 115 (2018) 900–906. 10.1016/j.ijbiomac.2018.04.138

[28] I. Mobasherpour, M.S. Heshajin, A. Kazemzadeh, M. Zakeri, Synthesis of nanocrystalline hydroxyapatite by using precipitation method, J. Alloys Compd. 430 (2007) 330–333. 10.1016/j.jallcom.2006.05.018.

[29] T. Hu, B. Cheng, Bone regeneration enhanced by green tea polyphenols/chitosan bifunctional hydrogel, Colloids and Surfaces B: Biointerfaces 114824 (2025). 10.1016/j.colsurfb.2025.114824.

[30] D. Liu, Z. Liu, J. Zou, L. Li, X. Sui, B. Wang, … B. Wang, Synthesis and characterization of a hydroxyapatite–sodium alginate–chitosan scaffold for bone regeneration, Front. Mater. 8 (2021) 648980. 10.3389/fmats.2021.648980.

[31] D.A. Gaber, H.S. Alhawas, F.A. Alfadhel, S.A. Abdoun, A.M. Alsubaiyel, R.M. Alsawi, Mini-tablets versus nanoparticles for controlling the release of amoxicillin: in vitro/in vivo study, Drug Des. Dev. Ther. (2020) 5405–5418. 10.2147/DDDT.S285522.

[32] S. Fanelli, A. Zimmermann, E.G. Totoli, H.R.N. Salgado, FTIR spectrophotometry as a green tool for quantitative analysis of drugs: practical application to amoxicillin, J. Chem. 2018 (2018) 3920810. 10.1155/2018/3920810.

